# Thermally tolerant symbionts may explain Caribbean octocoral resilience to heat stress

**DOI:** 10.1101/2020.10.21.349555

**Authors:** Jessie Pelosi, Katherine M. Eaton, Samantha Mychajliw, Casey P. terHorst, Mary Alice Coffroth

## Abstract

Coral reef ecosystems are under threat from the frequent and severe impacts of anthropogenic climate change, particularly rising sea surface temperatures. The effects of thermal stress may be ameliorated by adaptation and/or acclimation of the host, symbiont, or holobiont (host + symbiont) to increased temperatures. We examined the role of the symbiont in promoting thermal tolerance of the holobiont, using *Antillogorgia bipinnata* (octocoral host) and *Breviolum antillogorgium* (symbiont) as a model system. We identified five distinct genotypes of *B. antillogorgium* from symbiont populations isolated from *Antillogorgia* colonies in the Florida Keys. Three symbiont genotypes were cultured and maintained at 26°C (ambient historical temperature) and two were cultured and maintained at 30°C (elevated historical temperature) for two years. Following culturing, we analyzed the growth rate and carrying capacity of each symbiont genotype at both ambient and elevated temperatures in culture (*in vitro*). All genotypes grew well at both temperatures, indicating thermal tolerance among these *B. antillogorgium* cultures. Prior culturing at the elevated temperature, however, did not result in increased thermal tolerance. We then inoculated juvenile *A. bipinnata* polyps with each of the five symbiont genotypes, and reared these polyps at both ambient and elevated temperatures (*in hospite* experiment). All genotypes were able to establish symbioses with polyps in both temperature treatments. Survivorship of polyps at 30°C was significantly lower than survivorship at 26°C, but all treatments had surviving polyps at 56 days post-infection, suggestive of broad thermal tolerance in *B. antillogorgium*, which may play a part in the increased resilience of Caribbean octocorals during heat stress events.

## Introduction

In recent years, anthropogenic climate change has severely impacted coral reefs, which comprise some of the most biodiverse and economically important ecosystems on Earth (Hoegh-Guldberg 2011; Hoegh-Guldberg et al. 2017). The success of reef ecosystems in oligotrophic tropical waters is due to their obligate endosymbioses with dinoflagellate algae in the family Symbiodiniaceae (Muscatine and Porter 1977). In this mutualistic relationship, the coral host relies on photosynthate produced by its intracellular symbionts to thrive (Muscatine and Cernichiari 1969; Muscatine and Porter 1977). In exchange for these metabolites, the symbiotic algae recover host nitrogenous waste. This relationship, which is vital to the survival of the reef ecosystem, is currently threatened by anthropogenic environmental changes.

Coral bleaching, or the loss of algal symbionts, occurs due to a variety of environmental stressors including changes in salinity, pH, light intensity, and, most commonly, water temperature (Hoegh-Guldberg 1999). Most reef ecosystems now routinely encounter temperatures at or above their thermal maximum each year (Hoegh-Guldberg et al. 2007; Eakin et al. 2009), with bleaching events becoming more frequent and severe (Heron et al. 2016; van Hooidonk et al. 2016; Hughes et al. 2018; Oliver et al. 2018). As ocean temperatures continue to increase, it is essential to understand if and how corals and their symbionts can respond to the present threat.

The holobiont (host + symbiont) can respond to rising seawater temperatures via changes in the symbiont community within a single host, with the basic premise that hosts associating with more thermally tolerant symbionts will be more likely to survive bleaching events (i.e., shuffling and/or switching, *sensu* Baker 2003). Although most coral species host a single dominant symbiont type (Goulet 2006; LaJeunesse 2020), many hosts also harbor background or cryptic symbionts that are often undetected by traditional techniques (Mieog et al. 2007; Correa et al. 2009; Silverstein et al. 2012). During thermal stress, a more thermally tolerant background symbiont can become the dominant symbiont type, potentially conferring thermal tolerance to the holobiont (Berkelmans and van Oppen 2006; Jones et al. 2008; LaJeunesse et al. 2009; Keshavmurthy et al. 2012; Silverstein et al. 2015). In the aforementioned studies, the background symbionts often differed at the genus level and were typically replaced by the previously dominant symbiont type once the stress had subsided.

Thermally tolerant holobionts may arise through adaptation of symbiont populations, even in the absence of evolutionary changes in the host. Large population sizes, short generation times, and high mutation rates could greatly augment the standing genetic variation within symbiont populations (van Oppen et al. 2011). With greater standing variation, natural selection is more likely to produce populations with greater thermal tolerance (van Oppen et al. 2011; Torda et al. 2017; Quigley et al. 2018; Buerger et al. 2020). Numerous studies have established that more thermally tolerant symbionts can be selected from within a single symbiont genus or species (Howells et al. 2011; Chakravarti et al. 2017; Buerger et al. 2020). By sampling symbionts in the genus *Cladocopium* (ITS type C1) from reefs with different temperature regimes, Howells et al. (2011) showed that distinct symbiont genotypes had different thermal tolerances. Symbionts from the warmer reef displayed increased performance and survival at elevated temperatures as compared to those from the cooler environment. Furthermore, juvenile hosts inoculated with symbionts from the warmer reef performed better (grew faster, had less bleaching, and had lower mortality) than those inoculated with symbionts from the cooler region.

Chakravarti et al. (2017) and Buerger et al. (2020) presented evidence for selection within a species by examining the adaptive capabilities of the symbionts within a monoclonal culture of *Cladocopium goreaui*. Symbionts selected for thermal tolerance (i.e., those maintained at 31°C for 2.5 to 4 years) performed better at 31°C than the original monoclonal culture that had not undergone this selection treatment. Juvenile polyps infected with symbiont cells from the elevated temperature grew at 27°C and 31°C, while polyps infected with the original culture showed negative growth at 31°C. Infection with some strains of thermally selected cells, however, did not reduce the rate or severity of bleaching compared to those infected with the original culture (Buerger et al. 2020). These studies establish that selection for thermal tolerance can occur over ecologically relevant time scales within at least some symbiont populations. Given sufficient standing genetic variation, this is likely the case for other species within Symbiodiniaceae and requires further inquiry (Chakravarti et al. 2017).

By starting with a single monoclonal culture, all genetic variation in the symbionts used by Chakravarti et al. (2017) and Buerger et al. (2020) likely arose from *de novo* mutation. Intraspecific markers have revealed numerous instances where host colonies harbor multiple genotypes within a symbiont species (Santos et al. 2003; Pettay and LaJeunesse 2007; Howells et al. 2009; Kirk et al. 2009), suggesting the possibility for selection on pre-existing genetic variation within *in hospite* populations, resulting from both mutation and recombination (Baums et al. 2014; Chi et al. 2014; Wilkinson et al. 2015; Thornhill et al. 2017). Indeed, many studies have found variation in thermal tolerance within a single symbiont species (Parkinson and Baums 2014; Karim et al. 2015; Díaz-Almeyda et al. 2017; Grégoire et al. 2017; Bayliss et al. 2019). For example, Bayliss et al. (2019) identified genetic and trait variation within and among *Breviolum* species isolated from the octocoral *Antillogorgia bipinnata*.

Here we ask if there are thermally tolerant genotypes present in a symbiont species within a single host colony. We utilized the octocoral *A. bipinnata* and its host-specialist endosymbiotic algae, *Breviolum antillogorgium*, as a model to study the role of symbiont genetic variation in the thermal response of the holobiont. We first selected for thermally tolerant symbionts from *in hospite* populations to capitalize on pre-existing genetic variation. We then examined symbiont growth at two temperatures with an *in vitro* experiment using these cultured isolates of *B. antillogorgium*. Subsequently, we examined whether thermal tolerance in the symbiont could be conferred to the host using an *in hospite* experiment. The presence of thermally tolerant symbionts within a population contained in a single host colony expands the potential for the holobiont to ameliorate the consequences of increased temperatures, potentially slowing the loss of these invaluable ecosystems.

## Methods

### Symbiont isolation

Although culturing of Symbiodiniaceae is highly selective and rarely recovers the symbionts dominant within the host (Santos et al. 2001; LaJeunesse 2002), *Breviolum antillogorgium* (ITS2-type B1) has been successfully cultured and associates specifically with octocorals in the genus *Antillogorgia* (Santos et al. 2004; Parkinson et al. 2015). Symbiont cells were isolated from sixteen adult *Antillogorgia* colonies collected from 12-18 m depth at sites in Elbow and Pickles Reefs, Florida Keys in September 2016 (Table 1). A sample of each host colony was preserved in 95% ethanol for comparison of dominant host symbionts with those isolated. An approximately 2 cm piece was sampled from each colony and homogenized in 2 mL of filtered seawater (FSW). Each homogenate was passed through a series of mesh filters (250, 120, 70 and 20 µm), followed by several 1 mL 0.22 μm FSW rinses for a final volume of 3-6 mL. Homogenates were spun at 800 rpm for five minutes. Supernatants were decanted, and the pellets were resuspended in 10 mL of FSW. This was repeated three additional times, followed by resuspending each pellet in 1 mL of f/2 media (Guillard and Ryther 1962). For each colony, 50 µL of the resultant cell suspension was added to six flasks containing 30 mL of f/2 media. Three flasks of the symbionts from each colony were maintained at 26°C, mimicking typical thermal conditions, and three were immediately placed at 30°C to select for thermally tolerant genotypes that can survive at an elevated temperature. An elevated temperature of 30°C was chosen based on the historical thermal regime of the collection site during November/December, when newly settled *A. bipinnata* polyps acquire symbionts (Williams and Miller 2015; Online Resource 1). Initial growth, observed as a small dot at bottom of the flask, was immediately transferred into a new flask with fresh media. This was repeated each time new growth was observed in any flask over the first month, resulting in 131 cultures. All cultures were transferred to fresh media monthly.

### Genotype identification

After two years (approximately 735-1084 asexual generations, assuming a doubling time of 1.01-1.97 days, based on growth rates from this study), total symbiont genomic DNA from 106 cultures and the host source colonies was extracted using the 2X CTAB method (Coffroth et al. 1992) and diluted to 5-10 ng μL^-1^, and the flanking region of the B7Sym15 microsatellite (Pettay and LaJeunesse 2007) was sequenced (TACGEN, Richmond, California, USA). Sequences were compared to available sequences on NCBI using BLASTn for symbiont species-level identification.

Cultures identified as *B. antillogorgium* (n=24) and samples from the source colonies of these cultures were amplified with five microsatellite loci following published protocols: CA 6.38 (Santos et al. 2003), GV_1C (Andras et al. 2009), B7Sym8, B7Sym34, and B7Sym36 (Pettay and LaJeunesse 2007). Amplicons were labeled with fluorescently tagged primers and visualized on a 6.5% polyacrylamide gel with the LI-COR NEN Global IR2 DNA Sequencer (LI-COR Biotechnology Division) following Santos et al. (2003). Allele sizes were scored by eye relative to size standards between 50 and 350 bp. Based on the microsatellite results, 21 *B. antillogorgium* cultures were unialgal and contained one of five genotypes. Ultimately, this resulted in the occurrence of three distinct genotypes (referred to here as G1, G2, and G3) from one or more of the cultures reared at 26°C and two genotypes (referred to here as G4 and G5) from one or more of the cultures reared at 30°C (see Results). One of each of these five genotypes were used in the experiments outlined below (Table 1).

### *In vitro* growth experiments

Six replicates of each of the five cultures identified above were started by inoculating 18 mL of f/2 media with 10,000 cells mL^-1^. Three of the replicates for each culture were subsequently grown at 25.53 ± 0.89°C (mean ± SD; referred to as “26°C”) and three were grown at 29.84 ± 0.48°C (mean ±SD; referred to as “30°C”) on a 12:12 h light:dark cycle. Light levels were maintained at 120.58 ± 11.14 µmols m^-2^s^-1^ (mean ± SD) (Philips Energy Advantage T5 HO 49W) and 103.79 ± 37.3 µmols m^-2^s^-1^ (mean ± SD) (Philips T5 HO 24W/840) for the 26°C and 30°C incubators, respectively (Welch’s Two Sample t-test, p = 0.13). Cell densities were measured using six replicate hemocytometer cell counts every three days. Experiments were continued until cell density peaked and exhibited a consistent decline in abundance. We used the ‘growthrates’ package ver. 0.8.1 (Petzoldt 2019) in R ver. 4.0.2 (R Core Team 2020) to fit logistic growth curves to the cell counts over time, up to the maximum cell count for each culture. We extracted values of r and K from the ‘growthrates’ package and used these as estimates of maximum population growth rate and carrying capacity, respectively. Samples of each replicate were then preserved in 95% ethanol, from which DNA was extracted and amplified following the methods detailed above to verify symbiont genotype at the end of the experiment.

### *In hospite* polyp experiments

Branches from adult colonies of *A. bipinnata*, a gonochoric surface brooder (Kahng et al. 2011), were collected from Tennessee Reef, Florida Keys in November 2018 (N 24° 45.150’ W 81° 45.275’). These branches were transported to flow-through seawater tanks at Keys Marine Laboratory, where they were monitored daily for release of embryos. Aposymbiotic brooded larvae were collected from the colony surface on 16, 17, 22, 23, 25, and 26 November, washed three times in 0.22 μm filtered instant ocean (FIO), and transferred to Buffalo, NY. Following transport, larvae (4-14 days old) were again washed in FIO to remove environmental contaminants and groups of 25 larvae were transferred to pre-conditioned polypropylene containers (volume = approximately 115 mL) with 50 mL of FIO. For pre-conditioning, the bottom and sides of each container were roughened with sandpaper and then soaked for four weeks in 1.2 μm filtered seawater (FSW) to establish a biofilm. Once larvae were added, water in the containers (FIO) was changed daily for one week, then every other day until larvae settled and metamorphosed into polyps (approximately 4 weeks). Containers of newly settled polyps (approximately 1-2 weeks post-settlement) were then divided into five treatments (G1 through G5, Online Resource 2) and each treatment was inoculated with 1000 cells mL^-1^ of one of the five genotypes of *B. antillogorgium* utilized in the *in vitro* experiments (Table 1). For each genotype, 9-10 containers were placed at 26ºC (25.51 ± 1.10°C; mean ± SD) and 6-8 at 30ºC (29.79 ± 0.28°C; mean ± SD) and maintained in a 12:12 h light:dark cycle. Initial inoculations occurred twice over a period of one week. Polyp locations in the containers were mapped and survival for each individual polyp was recorded every 3-4 days with water changes. Visual infection, indicated by a light brown coloration in the polyp, was limited within the first month following initial introduction of symbionts. To increase the likelihood of the establishment of the symbiosis, inoculations were repeated with each water change during the second month and combined with feeding with Reefs-Roids (Polyplab, Quebec, Canada) (2 mL of Reef-Roids solution at a concentration of 1 mg/mL FIO) to induce symbiont uptake. Inoculations continued throughout the second month, and establishment of the symbiosis through visual observation was recorded throughout the experiment. An unidentified green alga, first observed in three containers on Day 42, but subsequently found in several additional containers on Days 56-59, led to extensive polyp mortality over time. After 69 days, all surviving polyps were collected, symbiont cell density was determined (as detailed above), and tissue was preserved in 95% ethanol. Symbiont DNA was extracted from preserved tissue and symbiont genotype was verified using the five microsatellite loci described above.

### Statistical analyses

We analyzed the effects of temperature treatment and genotype on *in vitro* symbiont maximum population growth rates and carrying capacity using separate two-way ANOVAs for each dependent variable (r and K), including the interaction term between these factors (temperature and genotype). Data were log-transformed to meet assumptions of homoscedasticity for the *in vitro* analyses. For the *in hospite* experiment, we quantified polyp survivorship as the proportion of surviving polyps after 31 and 52 days post-initial inoculation. These time points were prior to the observation of widespread contamination by the unidentified green alga, to avoid the possibility that this affected survivorship. Containers in which the contaminating algae was observed prior to Day 52 (n=5) were excluded from survivorship analyses beginning at the first observation of contamination. We analyzed survivorship over the first 31 days and then from Day 32 to Day 52 as described above; with survivorship for Days 1- 31 arcsine transformed to meet assumptions of normality (survivorship data for Days 32-52 were not transformed as the raw data fit the assumptions of the model without transformation). We analyzed temperature and genotype effects on visually observed infection at Days 31 and 52 as well as on symbiont cell densities at the end of the experiment (Day 69) as above. Symbiont cell densities were square-root transformed to meet assumptions of the model. Assumptions of normality and homogeneity were checked using residuals and Q-Q probability plots. All post-hoc pairwise comparisons were carried out using Tukey’s HSD (honest significance test) (Tukey 1949). *In vitro* statistical analyses were conducted in R ver. 4.0.2 and *in hospite* analyses in R ver. 3.6.1 (R Core Team 2020). R scripts are available at www.github.com/jessiepelosi/symbiont_tolerance and raw data are available at https://www.bco-dmo.org/award/658940.

## Results

### Sequencing and genotyping

Following two years of growth at either 26ºC or 30ºC, there were 131 viable cultures. The majority of cultures (106 of 131) either could not be classified or aligned most closely with *B. minutum*, a species commonly recovered from a range of species where it is undetected by normal techniques (LaJeunesse et al. 2012; Parkinson et al. 2015). Twenty-four of the cultures were identified as *B. antillogorgium* based on a comparison of B7Sym15 microsatellite flanking region sequences to available sequences on NCBI. Three of these cultures contained multiple genotypes. Each of the remaining 21 *B. antillogorgium* cultures were unialgal and contained one of five genotypes. Ten *B. antillogorgium* cultures reared at 26ºC were either genotype G1, G2 or G3, while 11 *B. antillogorgium* cultures reared at 30ºC were genotype G4 or G5. These five genotypes were used in the *in vitro* and *in hospite* experiments. Based on microsatellite genotyping, one of the five genotypes (G3) was the same as the dominant genotype in the source colony; the other four genotypes represented background symbionts as is common in culturing from host tissue (Santos et al. 2001; LaJeunesse 2002; LaJeunesse et al. 2012; Parkinson et al. 2015).

### *In vitro* growth experiments

Cultures reached their maximum cell density in 28-40 days and maintained this density for at least several days afterward (Fig 2a, Online Resource 3). The effect of temperature on maximum population growth rate was dependent on the symbiont genotype (Temperature*Genotype: F_4,20_ = 30.20, p < 0.001). At 30ºC the mean growth rate for two of the five genotypes (G1 and G2) was significantly lower than that at 26ºC; there was no significant effect of temperature for the remaining three genotypes (Fig. 2a, b). The effect of temperature on carrying capacity also depended on the symbiont genotype (Temperature*Genotype: F_4,20_ = 17.90, p < 0.001). One genotype (G3) showed a decrease in carrying capacity at the elevated temperature, but the remaining genotypes showed non-significant responses to temperature (Fig. 2c). The time for populations to reach their peak abundance was also dependent on the interaction between temperature and symbiont genotype (Temperature*Genotype: F_4,20_ = 6.57, p = 0.002).

**Figure 1.**
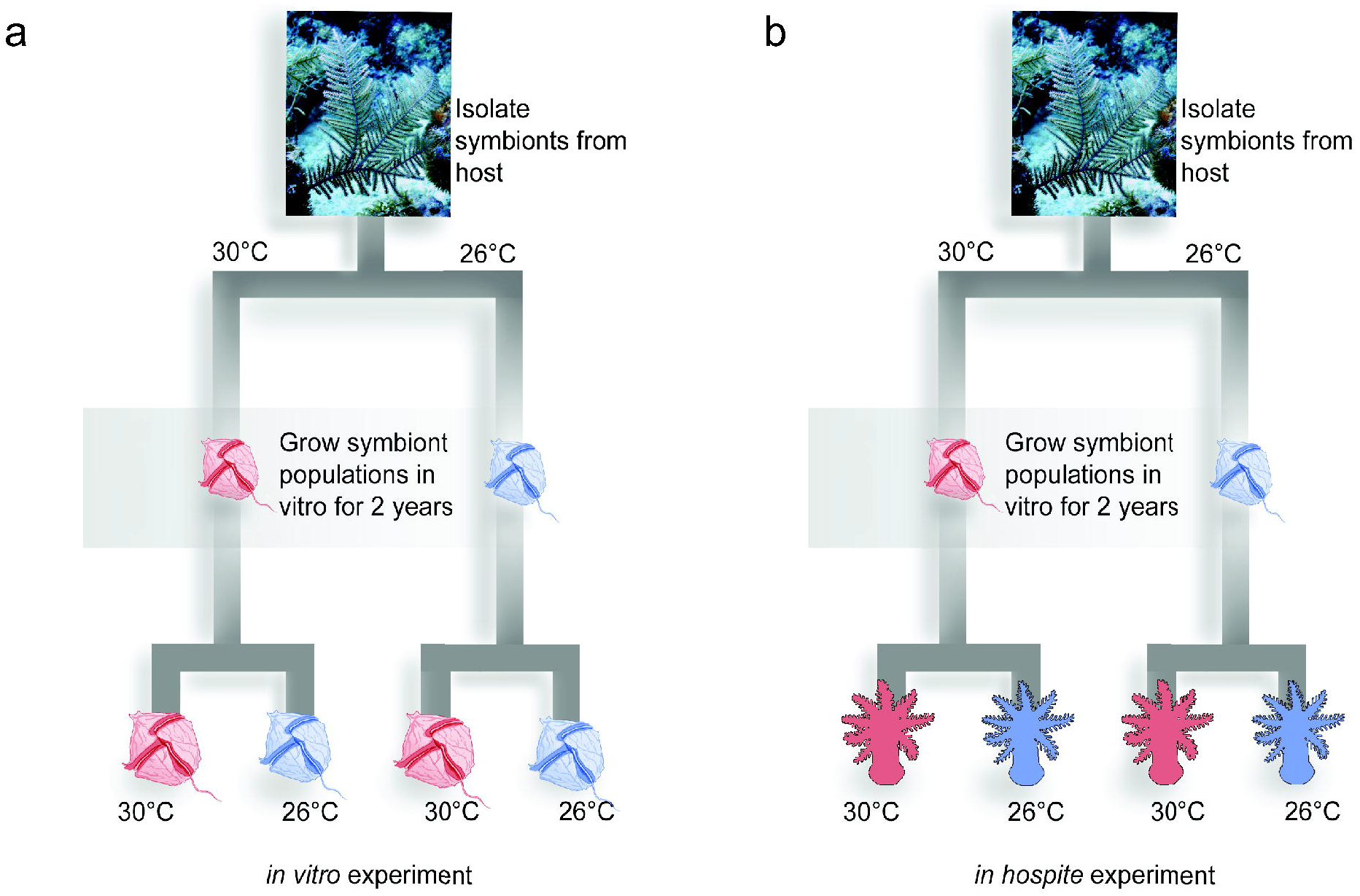
Experimental design for a) *in vitro* growth experiment and b) *in hospite* experiment. Figure colors refer to historical temperature treatment (blue, maintained at 26°C; red, maintained at 30°C), with tip figures colored based on experimental treatment (not historical temperature).

**Figure 2.**
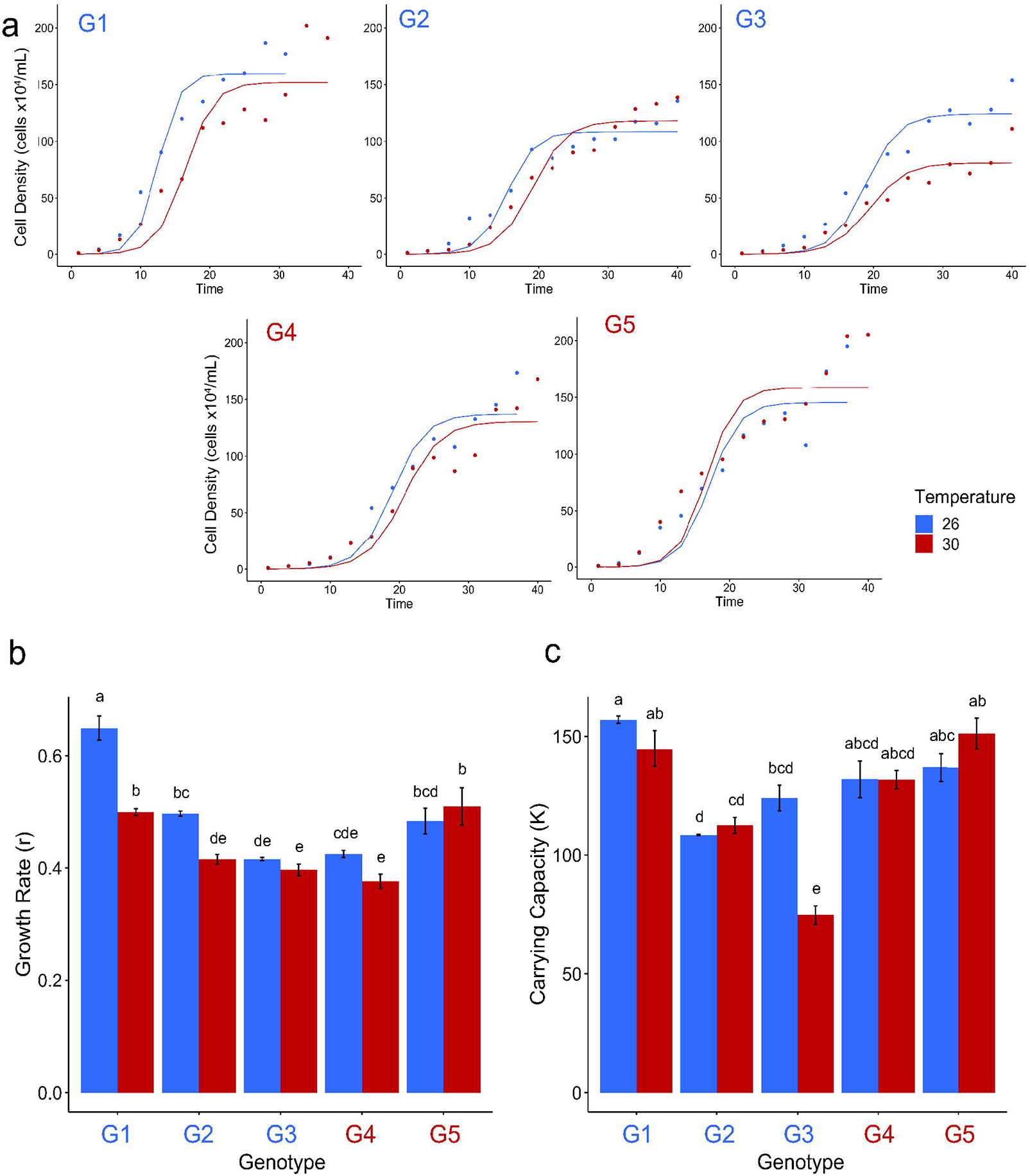
a) Mean logistic growth fits to *in vitro* cell densities of each genotype of *Breviolum antillogorgium* over time. Blue represents growth at 26°C and red at 30°C. b) Estimated mean (+/- SE) maximum symbiont population growth rates (r) and c) estimated mean (+/- SE) carrying capacities (K). Colors for genotype names (x-axis) refer to the historical temperature at which they were grown, blue indicating cultures maintained at 26°C and red indicating cultures maintained at 30°C two years prior to the start of the experiment. Letters above bars represent differences among groups based on Tukey post-hoc tests.

### *In hospite* polyp experiments

For the first 31 days, we did not find a significant impact of genotype (F_4,73_ = 1.84, p = 0.130), temperature (F_1,73_ = 0.03, p = 0.861), nor Temperature*Genotype (F_4,73_ = 0.83, p = 0.508) on the proportion of visually infected polyps (Fig. 3a). Similarly, between 32 and 52 days, we did not find a significant impact of genotype (F_4,56_ = 0.66, p = 0.623), temperature (F_1,56_= 3.77, p = 0.057), nor Temperature*Genotype (F_4,56_ = 1.22, p = 0.312) on the proportion of visually infected polyps (Fig. 3b). We found a significant interaction between the effects of temperature and genotype on the symbiont cell densities in polyps at the end of the experiment, when the vast majority of polyps (range 66-100% of polyps per treatment) were infected (69 days post first inoculation, F_4,264_ = 6.34, p < 0.001). Average symbiont densities in polyps varied between genotypes, depending on temperature at Day 69 (termination of the experiment, Fig. 3c). Polyp survivorship at the end of the first 31 days was not significantly affected by symbiont genotype (F_4,74_ = 0.81, p = 0.522), nor by treatment temperature (F_1,74_ = 3.05, p = 0.085), and there was not a significant interaction term (F_4,74_ = 0.76, p = 0.574) (Fig. 4a, b). Polyp survivorship at the end of 52 days decreased dramatically in both temperature treatments (Fig. 4a, c), with those polyps at 30°C having significantly lower survivorship than those reared at 26°C (F_1,68_ = 15.64, p < 0.001). There was not a significant effect of genotype (F_4,68_ = 2.11, p = 0.089) nor the interaction between temperature and genotype on survival for the second period (Temperature*Genotype, F_4,68_ = 0.13, p = 0.970). Including the five containers removed after Day 42 (due to algal contamination) did not change the results. Although survivorship results are only reported prior to the appearance of an unidentified green alga, all treatments were maintained for 69 days. At that time, all treatments except G4 at 30ºC had surviving polyps (range 2-70 polyps per treatment).

**Figure 3.**
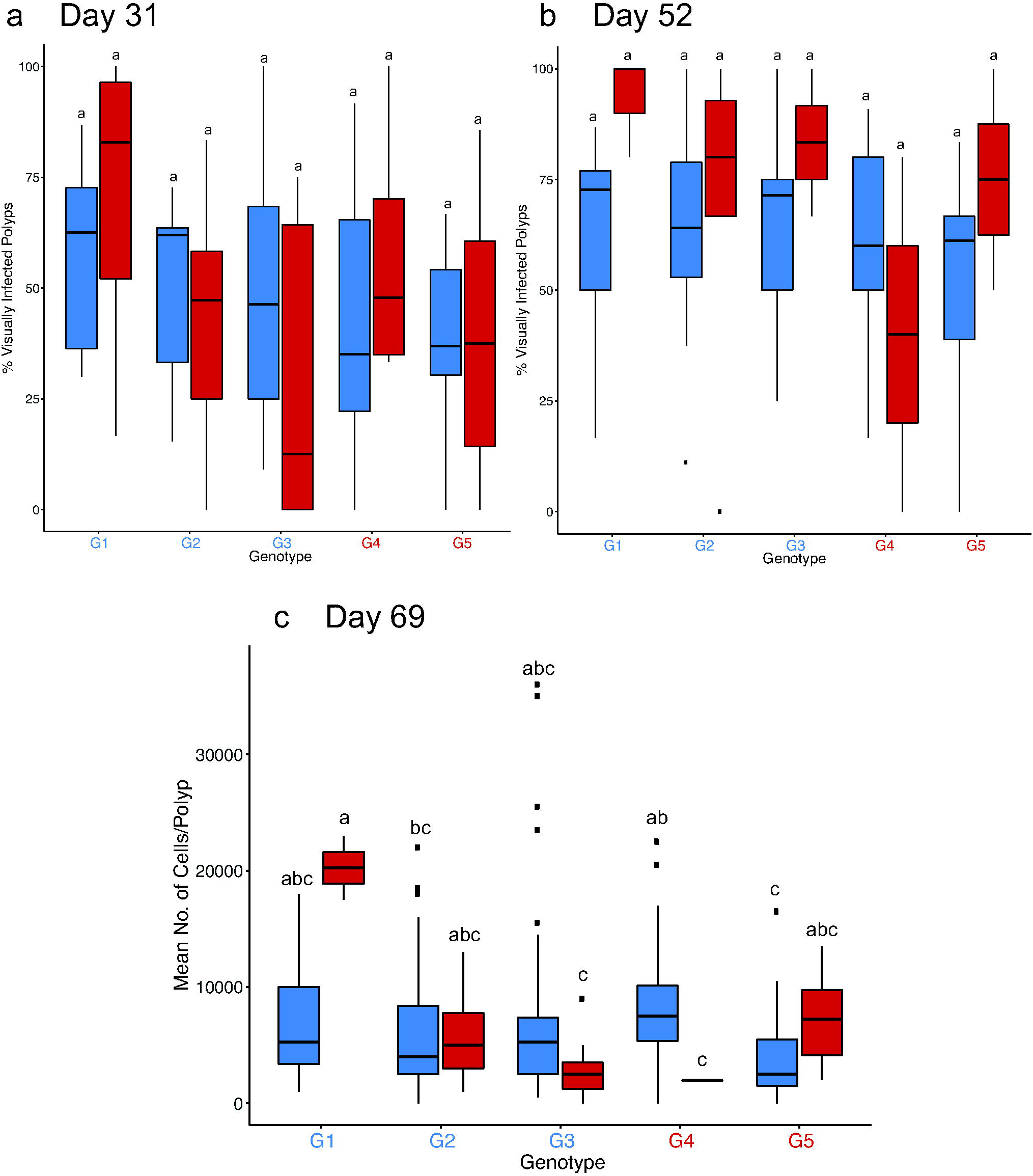
Mean percent of visually infected polyps at the end of a) Day 31 and b) Day 52. c) Symbiont cell densities within *Antillogorgia bipinnata* polyps inoculated with five symbiont genotypes at 26ºC and 30ºC at the end of the experiment (Day 69). Colors for genotype names (x-axis) refers to historical temperature, blue indicating cultures maintained at 26ºC and red indicating cultures maintained at 30ºC two years prior to the start of the experiment. Color of boxes refers to experimental temperature (blue-26ºC, red-30ºC). Boxes represent interquartile range and whiskers the range of the data. Points indicate outliers. Letters above boxes represent differences among groups based on Tukey post-hoc tests.

**Figure 4.**
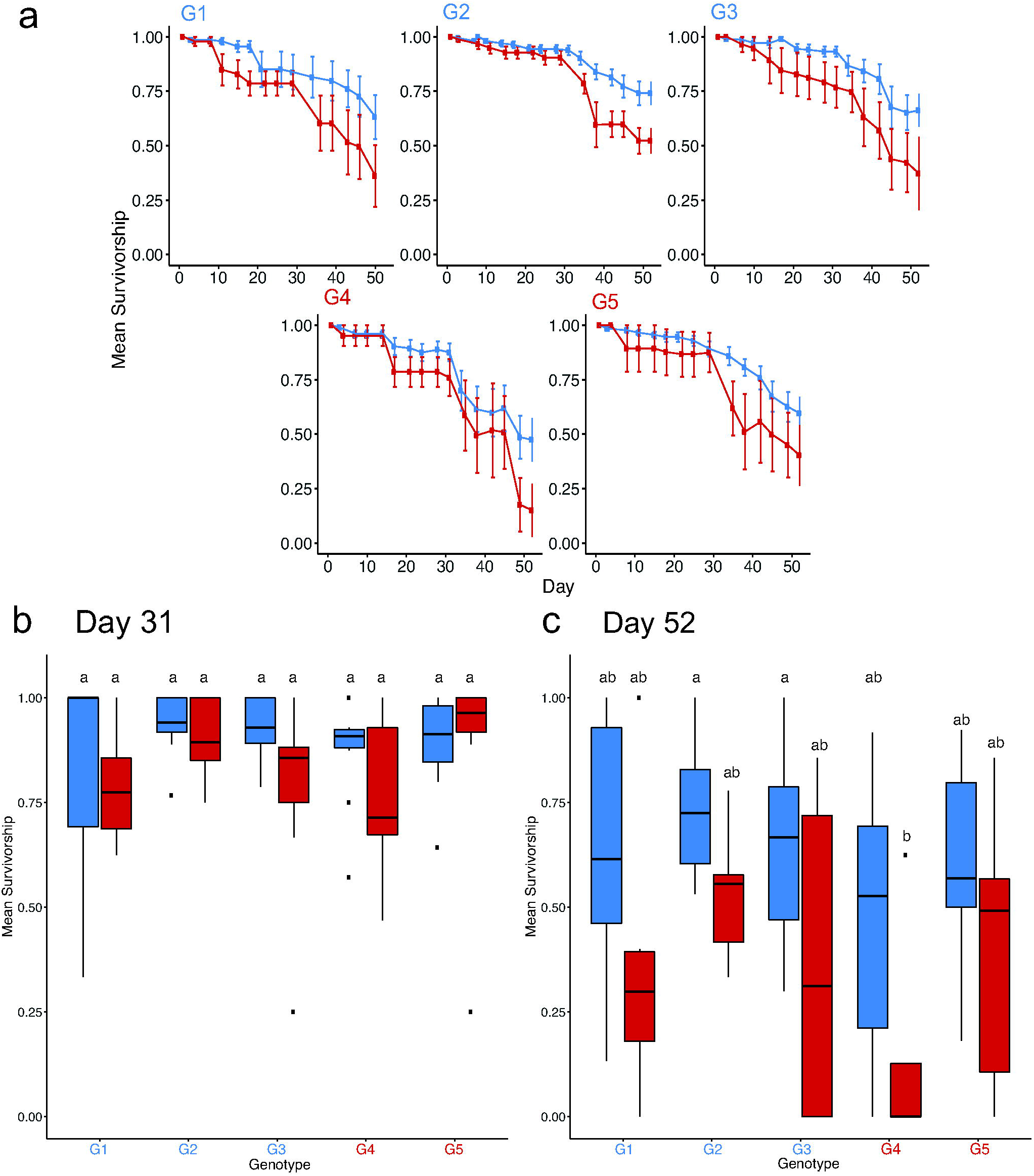
a) Mean polyp survivorship over time for *A. bipinnata* polyps inoculated with five symbiont genotypes at 26ºC and 30ºC. Error bars are ±SE. Blue lines refer to treatments at 26 ºC and red lines refer to treatments at 30ºC. Differences in average survivorship at b) Day 31 and c) Day 52. Colors for genotype names (x-axis) refer to historical temperature, blue indicating cultures maintained at 26ºC and red indicating cultures maintained at 30ºC two years prior to the start of the experiment. Color of boxes refers to experimental temperature (blue-26ºC, red-30ºC). Boxes represent interquartile range and whiskers the range of the data. Points indicate outliers. Letters above boxes represent differences among groups based on Tukey post-hoc tests.

## Discussion

Cultured symbionts isolated from standing genetic variation within host colonies revealed the presence of multiple genotypes capable of growth at 30ºC, with three of the five unique genotypes used here isolated from a single host colony (Table 1). However, there was no clear evidence of selection for increased thermal tolerance in cultures reared at 30ºC for over 700 generations (2 years) compared to those reared at 26ºC over this time (Figs. 2-4). Genotypic variability was clear as genotypes differed in their response to thermal stress (i.e., growth rate and carrying capacity, Fig. 2). Regardless of this variability, all tested genotypes showed growth at 30ºC, suggesting that *Breviolum antillogorgium*, as a species, is thermally tolerant.

### *In vitro* growth experiments

By isolating genotypes that were able to grow at elevated temperatures from standing genetic variation among *in hospite* populations, we established that thermally tolerant symbiont genotypes are present within the host. We maintained these cultures at 30°C for two years to select for long-term growth at elevated temperatures, anticipating that these cultures would adapt and outperform other genotypes at elevated temperatures. Unlike previous studies that found evidence for thermal adaptation in Symbiodiniaceae after one to 2.5 years of exposure to increased temperatures (*Cladocopium* - Chakravarti et al. 2017; *Fugacium, Gerakladium, Symbiodinium -* Chakravarti and van Oppen 2018), a history of long-term growth at 30°C did not yield better performance for *B. antillogorgium* at 30°C (as compared to 26°C). Our results suggest that local adaptation to this temperature did not occur over this time period. It is possible that all examined genotypes possessed pre-existing adaptations to temperatures at or above 30ºC, as over the past decade and a half, if not longer, this threshold has been frequently exceeded during the summer months in the Florida Keys (Online Resource 1).

Growth rate serves as a good proxy for fitness of symbiont populations. Carrying capacity may serve as a proxy for efficiency of resource use because it suggests that more organisms can be sustained at the same level of resource. We found variation among genotypes in carrying capacity, but interestingly, also found variation in the response of carrying capacity to increased temperature. One genotype (G3) showed a significant decrease in carrying capacity at higher temperature (Fig. 2c), but the remaining four genotypes showed non-significant responses to temperature. The ability of all cultures to survive and grow positively at 30°C for several months, regardless of historical temperature, suggests that these genotypes of *B. antillogorgium* possess pre-existing, broad thermal tolerance, possibly due to plastic acclimatory responses.

Prior studies examining the potential for symbionts to adapt to elevated temperatures have focused on a single genotype (Chakravarti et al. 2017; Chakravarti and van Oppen 2018), but in the present study we used several genotypes within the same species, selected from pre-existing genetic variation within *in hospite* populations. Variability in response to elevated temperatures observed among the genotypes used in our experiments highlights the potential role of intraspecific genetic variation in responding to thermal stress and further underscores the importance of examining both intrageneric and intraspecific variation in thermal responses. Other *in vitro* studies (Chakravarti et al. 2017; Chakravarti and van Oppen 2018; Buerger et al. 2020) have explored how dinoflagellate symbionts in other genera respond to temperatures higher (31°C-34°C) than used in our experiments. It is possible that the rapid adaptation seen in previous studies might also occur in *B. antillogorgium* if these genotypes were grown at higher temperatures. It is clear that elevated temperatures in excess of 30ºC are becoming frequent during the summer months in the Florida Keys (Online Resource 1) and further work should explore the limits of thermal tolerance in *B. antillogorgium* to temperatures greater than 30°C.

### *In hospite* polyp experiments

There were no significant differences in the percent of visually infected polyps between genotypes and temperature treatments through Day 52. In general, the percent of infected polyps on Day 52 was largely congruent with the average number of symbiont cells per polyp at the end of the experiment; those genotypes that had a greater proportion of visibly infected polyps also had a greater average number of symbiont cells per polyp (Fig. 3b, c). Final symbiont cell density varied with genotype and temperature, indicating that symbionts were either taken up or grew at different rates within hosts. The lack of a consistent effect of temperature on symbiont density within polyps suggests that the establishment of the symbiosis at 30°C may be as stable as those at ambient temperatures (26°C). Given the significant interaction between treatment temperature and symbiont genotype on symbiont density within polyps at the end of the experiment (Day 69), the ability of the symbiont to infect and thrive in the host likely depends on multiple factors, with potential for variation in the establishment and maintenance of the symbiosis (Davy et al. 2012; Hawkins et al. 2016). It is clear that biotic interactions (e.g., recognition and phagocytosis of algal symbionts, Davy et al. 2012) drive the establishment, while abiotic factors, such as temperature, may dictate the maintenance of the symbiosis (e.g., Rädecker et al. 2021).

The overall thermal tolerance recorded *in vitro* was observed in the polyp experiment, even though performance *in vitro* was not directly mirrored *in hospite* (i.e., those symbiont genotypes that had the highest growth rates *in vitro* did not necessarily confer the highest survivorship under the same conditions *in hospite*). All treatments had relatively high polyp survival early in the experiment, with a more rapid decline in survivorship towards the end (Fig. 4). Polyps showed sustained survivorship over an extended period of heat exposure, with some surviving polyps in all treatments at 56 days post-inoculation. Furthermore, after 69 days, all treatments except G4 at 30ºC had surviving polyps. This is a substantially longer exposure to heat stress than has been previously reported for *in hospite* experiments (i.e., Chakravarti et al. 2017: 28 days; Buerger et al. 2020: 7 days). Our results suggest that the observed thermal tolerance exhibited by our symbiont types may have contributed to or enhanced the thermal tolerance of the holobiont, although the long-term value of this has yet to be determined.

In this study, the host-symbiont pairs exhibited thermal tolerance. At the elevated temperature, however, hosts infected with *Breviolum* cultures that had been reared at 30°C did not consistently outperform their counterparts (i.e., hosts infected with cultures reared at 26°C). This indicates that selection for thermal tolerance of certain genotypes likely occurred prior to the isolation of these symbionts. Increasingly elevated temperatures are becoming more commonplace on coral reefs (Heron et al. 2016; van Hooidonk et al. 2016; Hughes et al. 2018; Oliver et al. 2018), and repeated exposure to these elevated temperatures may have selected for symbiont genotypes capable of surviving such extremes. Ultimately, our results indicate that there exists standing genetic variation within symbiont populations, which in this case, contained genotypes that were thermally tolerant. These symbionts may be vital for the holobiont’s survival, although the basis for this thermal tolerance requires further investigation.

### Role of the symbiont and implications for Caribbean reefs

Most previous studies of the capacity for rapid thermal adaptation in symbionts have focused on Pacific strains of Symbiodiniaceae. Our results shed new light on the existence of thermally tolerant Caribbean coral endosymbionts in genera in addition to those thermally tolerant species within the genus *Durusdinium* (LaJeunesse et al. 2014). We provide key evidence that even in the absence of rapid adaptation to elevated temperatures, thermal tolerance may be widespread in some *Breviolum* species. It is likely that broad thermal tolerance is an inherent trait among these genotypes and perhaps among other species within *Breviolum*, particularly those within the B1 (ITS2-type) lineage (Pleistocene (B1) radiation, *sensu* LaJeunesse 2005; Parkinson et al. 2015) such as *B. antillogorgium*. Under induced bleaching and/or thermal stress, for example, the octocoral *Briareum asbestinum* lost its dominant *Breviolum* ITS2 type B19 symbiont, which was replaced by a *Breviolum* ITS2 type B1 species (Lewis and Coffroth 2004; Poland and Coffroth 2019). Transitioning from a dominant symbiont species with lower thermal tolerance to a more thermally tolerant cryptic species during bleaching events may buffer Caribbean octocorals from increasing temperatures.

Coral reefs are in decline worldwide and some regions, such as the Caribbean, are experiencing a phase shift from scleractinian-dominated to octocoral-dominated ecosystems (Ruzicka et al. 2013; Lenz et al. 2015; Lasker et al. 2020). This pattern has been attributed, in part, to high levels of octocoral recruitment (Lasker et al. 2020). However, bleaching in octocorals is rarely observed during mass bleaching events (Lasker et al. 1984; Lasker 2003; Prada et al. 2010) and if it is observed, most species show a high level of resilience (Prada et al. 2010). The fact that species within *Breviolum* are the dominant symbionts in Caribbean octocorals (LaJeunesse 2002; van Oppen et al. 2005; Goulet et al. 2008), and that some symbiont species within this genus appear to be thermally tolerant, as shown here, may contribute to this resilience. Additional studies are needed to determine the prevalence of these thermally tolerant symbionts, which may exist at background level, and their role in mitigating the effects of increasing temperatures on reef cnidarians. The survival of Caribbean reefs may be reliant on evolutionary rescue (Gomulkiewicz and Holt 1995) perpetrated by selection for these more thermally tolerant symbiont genotypes within a given species; a process that could be particularly important for specialist hosts that are not able to harbor different species of symbionts.

## Supporting information

Online Resource 1

Online Resource 2

Online Resource 3

## Acknowledgements

We would like to thank H. Franklin, undergraduate volunteers, staff of the KML, the FKNMS and the FWC (Permit FKNMS-2014-018-A2 and SAL-18-2052-SR, respectively). This work was supported by NSF-OCE-1559286 (MAC) and NSF-OCE-1559105 (CPt), a UB Honors Research & Creativity Grant (JAP), and a UB CURCA Award (KME). We are grateful to D. Williams for providing the NOAA temperature data and to H. Lasker, A. Martinez, C. Wells, and two anonymous reviewers for helpful and insightful comments on previous drafts of this manuscript.

**Online Resource 1**. Average water temperature at Elbow Reef A at 3 m depth from 2005 to 2015. Data from Williams and Miller (2015). Horizontal lines are at 30ºC (red) and 26ºC (blue) for reference.

**Online Resource 2**. Experimental set up for the *in hospite* study. Letter color refers to historical temperature, blue indicating cultures reared at 26ºC and red indicating cultures reared at 30ºC two years prior to the start of the experiment.

**Online Resource 3**. Mean logistic growth fits to *in vitro* cell densities of each genotype of *Breviolum antillogorgium* over time for each replicate for each genotype and temperature treatment.

