## Supplementary material for "Thermally tolerant symbionts may explain Caribbean octocoral resilience to heat stress": Table 1

**Table 1.** Five genotypes of *Breviolum antillogorgium* selected for *in vitro* and *in hospite* experiments. Letter color refers to selection temperature, blue indicating cultures maintained at 26°C and red indicating cultures maintained at 30°C two years prior to the start of the experiment. Cultures were isolated from *Antillogorgia* colonies collected at Elbow Reef, Florida Keys.

| Genotype | Selection Temp. | Source Colony | Reef | Depth | Location | Accession No. |
|---|---|---|---|---|---|---|
| G-1 | 26°C | 7 | Elbow | 18 m | N 25 07.925<br>W 80 15.717 | MW207288 |
| G-2 | 26°C | 5 | Elbow | 11 m | N 25 07.956<br>W 80 15.810 | MW207289 |
| G-3 | 26°C | 8 | Elbow | 18 m | N 25 07.925<br>W 80 15.717 | MW207287 |
| G-4 | 30°C | 8 | Elbow | 18 m | N 25 07.925<br>W 80 15.717 | MW207285 |
| G-5 | 30°C | 8 | Elbow | 18 m | N 25 07.925<br>W 80 15.717 | MW207286 |
