## Supplementary figures and images for "Thermally tolerant symbionts may explain Caribbean octocoral resilience to heat stress"

### Online Resource 1

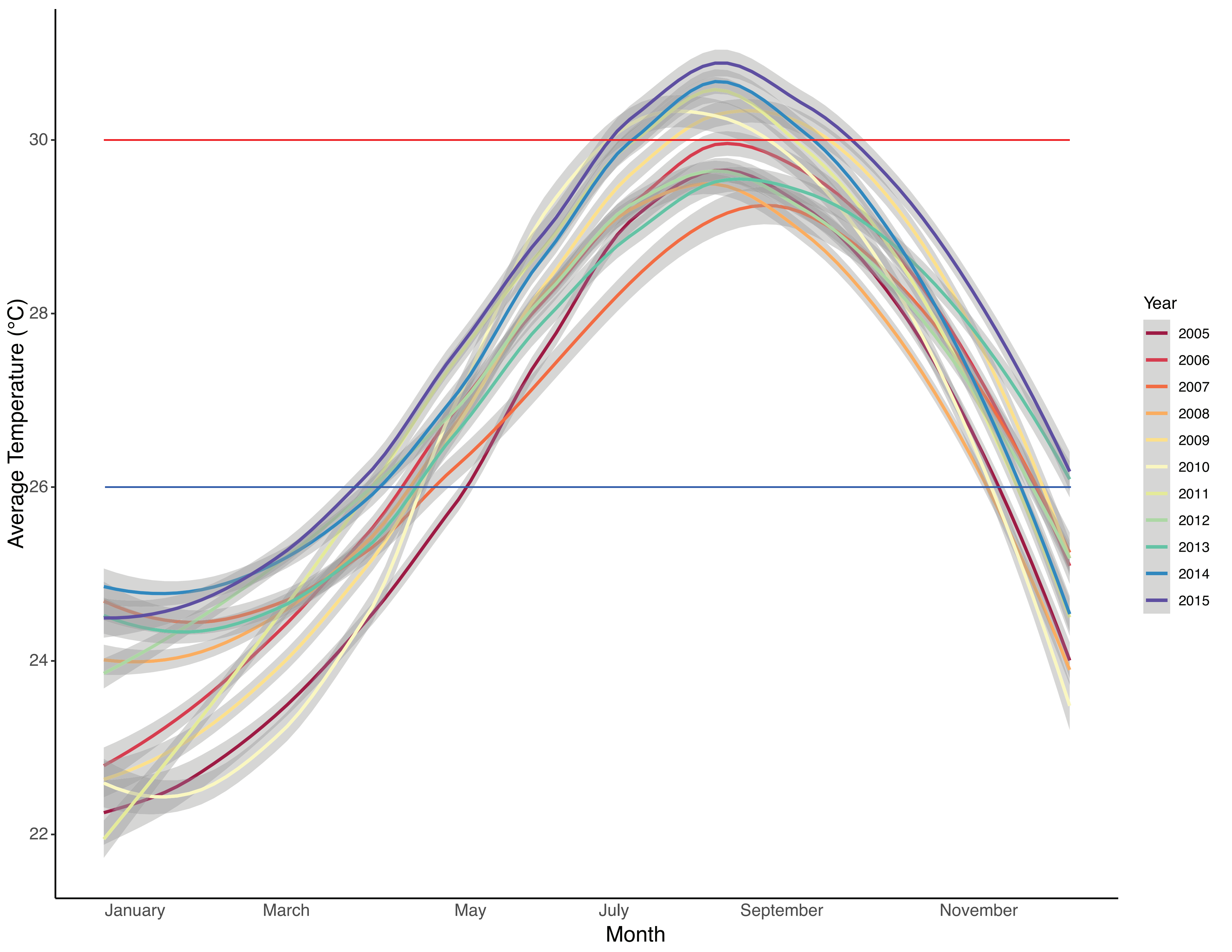

### Online Resource 3

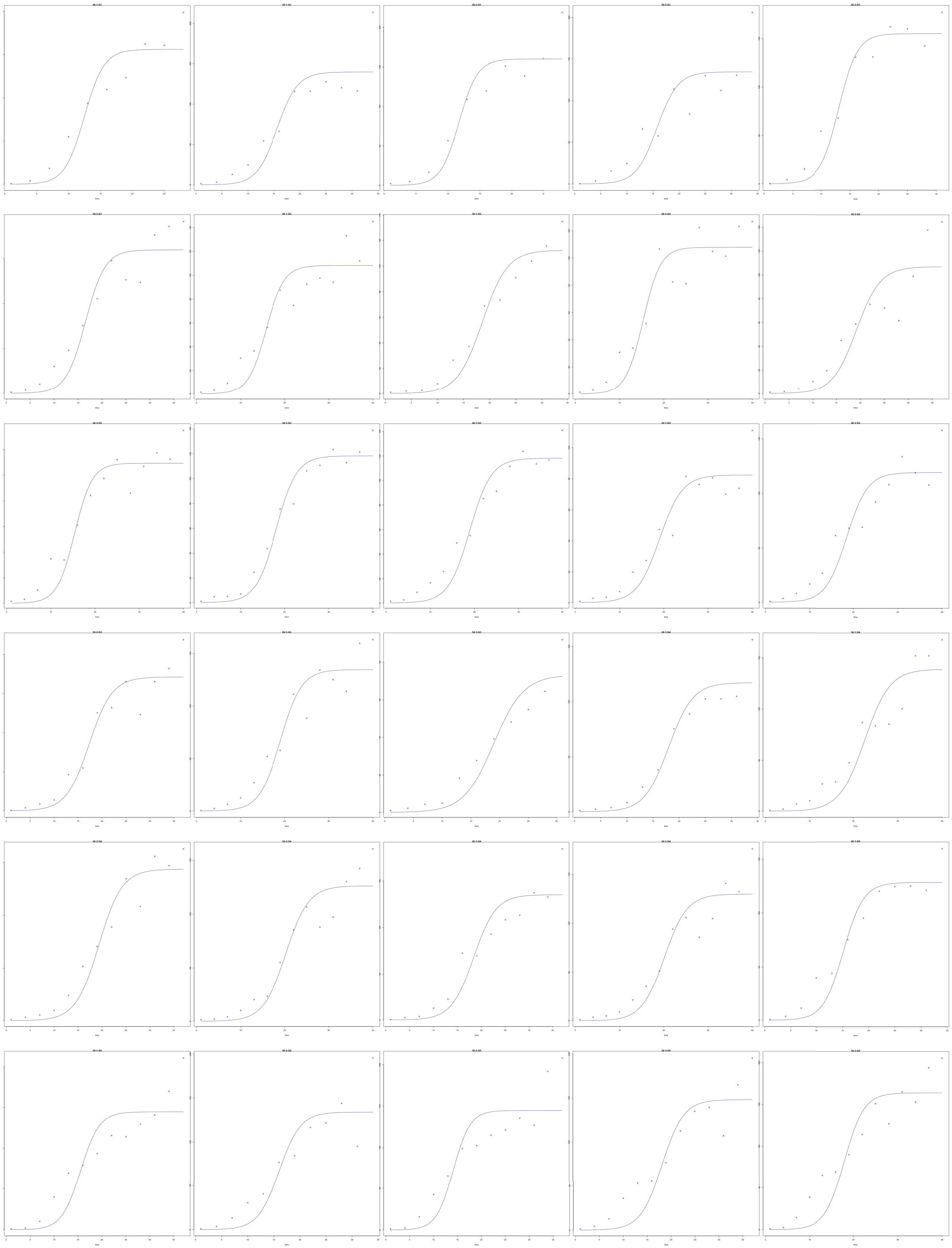
