## Supplementary material for "Thermally tolerant symbionts may explain Caribbean octocoral resilience to heat stress": Online Resource 2

**Online Resource 2.** Experimental set up for the *in hospite* study. Letter color refers to selection temperature, blue indicating cultures reared at 26ºC and red indicating cultures reared at 30ºC two years prior to the start of the experiment.

| Treatment | Mean # Polyps per Container | # Polyps per Treatment | Containers per Treatment |
|---|---|---|---|
| G-1 @ 26ºC | 13 | 117 | 9 |
| G-2 @ 26ºC | 14 | 135 | 10 |
| G-3 @ 26ºC | 14 | 136 | 10 |
| G-4 @ 26ºC | 13 | 133 | 10 |
| G-5 @ 26ºC | 13 | 132 | 10 |
| G-1 @ 30ºC | 8 | 46 | 6 |
| G-2 @ 30ºC | 9 | 71 | 8 |
| G-3 @ 30ºC | 8 | 54 | 7 |
| G-4 @ 30ºC | 8 | 49 | 6 |
| G-5 @ 30ºC | 9 | 61 | 7 |
